# Cyclohexyl acetate functions like a volatile sex pheromone mimic in *Caenorhabditis nematodes*

**DOI:** 10.1101/2025.09.19.677433

**Authors:** Xuan Wan, Yuki Togawa, Matthew R. Gronquist, Marika Sagawa, Daniel Leighton, Chung Man Chan, Frank C. Schroeder, King L. Chow, Paul W. Sternberg, Ryoji Shinya

## Abstract

Nematodes communicate via diverse sex pheromones, including long-range volatile signals, short-range chemical cues, and contact-dependent molecules. While the ascaroside family of small molecules that mediate short-range attraction is well characterized, the identities and roles of volatile sex pheromones (VSPs) that act over longer ranges remain unknown. Using GC-MS analysis of crude VSP extracts, we identified cyclohexyl acetate (CA) as a candidate mimic, sharing retention time and mass spectral features with natural VSPs. Behavioral assays demonstrated that CA acts as a concentration-dependent, male-specific attractant in *Caenorhabditis.* Pre-exposure to VSPs induced cross-adaptation to CA, suggesting shared sensory processing. Surprisingly, genetic and calcium imaging analyses revealed that CA perception is mediated primarily by AWC_on_ (*str-2*-expressing) neurons and involves VSPs chemoreceptor *srd-1*-independent pathways, which are distinct from the neural pathways involved in natural VSPs perception. These findings establish that CA is not a major component of VSPs but a structural and functional mimic of nematode VSPs, operating through a parallel sensory circuit. Although the endogenous source of CA remains unknown, its structural and behavioral mimicry provides new insights into the complexity of chemosensory signaling and the potential for interspecies chemical eavesdropping in nematode ecology.

## Introduction

Communication via chemical cues is ubiquitous across the animal kingdom. Small molecule chemical signals mediate a variety of intraspecific and interspecific interactions, such as mate location and choice and predator-prey encounters. In the nematode *Caenorhabditis elegans*, a family of small molecules, the ascarosides, controls sexual attraction, development, and social behavior [1, 2]. Blends of these compounds, dauer pheromones, integrate environmental cues like population density and food availability to trigger larval entry into the stress-resistant dauer stage, a key survival adaptation under adverse conditions [3, 4]. Beyond developmental regulation, ascarosides influence behaviors ranging from mate attraction and aggregation to repulsion and exploratory movement [2, 5]. Some exhibit concentration-dependent function; for example, ascr#2 and ascr#3 attract males at low concentrations but repel hermaphrodites at higher concentrations [4, 5]. This functional versatility shows the complicated context-dependency of pheromone communication [6-11].

While nonvolatile pheromones such as ascarosides mediate contact-dependent, short-range communication, *C. elegans* also utilizes volatile sex pheromones (VSPs) for long-range mate attraction. VSPs can attract males from distances of at least 10 cm under laboratory conditions [12]. Although ascarosides also can diffuse through the environment, they are generally classified as nonvolatile and require direct contact or proximity to trigger behavioral responses [2, 5, 13]. Among ascarosides, only certain mate-searching compounds such as ascr#2, ascr#3, and ascr#4 are active over relatively longer ranges [2, 14].

The VSP facilitates mate attraction and provides critical information about an individual’s sex, reproductive status, and developmental stage [12]. The existence of long-range volatile signals is supported by observations that certain chemical communications persist even in mutants deficient in ascaroside production [15, 16]. Virgin females from the *Caenorhabditis* species *C. elegans, C. remanei,* and *C. brenneri* secrete VSPs that is attractive to males from the closely related androdioecious species *C. elegans and C. briggsae*, as well as the dioecious *C. brenneri* and *C. remanei*. The production of VSPs is both sex- and stage-specific. In *C. elegans* hermaphrodites, oocyte–somatic communication regulates the synthesis of mate-attracting VSPs. These volatiles are produced only by virgin *fog-2* mutant females or by hermaphrodites that have exhausted their self-sperm [12, 15].

Sexual attraction in *C. elegans* is mediated by a combination of sex-specific and core sensory neurons. Male-specific neurons, such as CEMs, and sex-shared sensory neurons, such as AWAs, contribute to robust male attraction [15, 17-19]. The male response to volatile cues relies on molecular pathways involving G protein-coupled receptors and TRPV channels, such as OSM-9 [15, 17, 18]. Moreover, the sexually dimorphic expression of receptors, such as SRD-1 only in the AWA neurons of males, enhances male sensitivity to VSP signals, reflecting the nuanced regulation of chemosensory signaling [18].

Chemical signaling is highly conserved in nematodes [20], and other species have evolved the ability to detect and respond to these signals [21, 22]. The influence of *C. elegans* pheromones extends beyond nematodes, affecting interkingdom interactions. For example, nematophagous fungi, which are natural predators of soil-dwelling nematodes, can detect and respond to nematode-secreted ascarosides to induce trap formation, a coevolved predator-prey relationship [23]. Similarly, plants, including *Arabidopsis*, tomato, potato, and barley, respond to ascarosides such as ascr#18 by activating robust defense mechanisms against a wide range of pathogens, including viruses, bacteria, and fungi [24]. Plant-parasitic nematodes utilize ascarosides to promote the reproductive development of their vector beetles, which, in turn, secrete ascarosides to attract nematode larvae for dispersal [25].

The lack of clarity regarding the identities of VSPs produced by *C. elegans* and related species represents a significant gap in our understanding of communication within nematodes. While over 200 ascaroside variants have been characterized [26], the identities of VSPs remain elusive despite close to two decades of research and the identification of numerous candidate compounds. To address this gap, we conducted a targeted screen of potential VSP analogs. Here, we report the identification of cyclohexyl acetate (CA) as a chemical mimic of the volatile, long-range, male-attractive pheromone found among *Caenorhabditis* nematodes.

## Methods

### Volatile Compound Collection for GC-MS

The mated *C. remanei* (wild-type EM464) female loses the ability to produce VSPs [12]. Males and female can be reliably distinguished based on morphology beginning in the L4 larval stage. To obtain virgin female for experiments, L4-stage animals were isolated from mixed populations. The operational window for virgin female collection spans approximately 15–16 hours. To extend this period and obtain sufficient numbers, L1-arrested worms were released at three time points spaced 2 hours apart, expanding the collection window to 20 hours. The following day, virgin females were confirmed by the absence of eggs on the plate.

A total of 5,000 one-day-old *C. remanei* were collected for each sample and washed three times with M9 buffer. Volatile components were isolated using a purge-and-trap system with a Tenax TA adsorbent liner (poly(2,6-diphenyl-p-phenylene oxide), 60–80 mesh [27]. Tenax TA selectively traps volatile and semi-volatile analytes within a molecular range of n-C₇ to n-C₂₆. As illustrated in Figure S1, samples (2 mL) were loaded into a glass vial sealed with a rubber stopper and two gas-flow needles. The vial was placed in a heated water bath (50 °C), and volatile compounds were purged under a nitrogen stream (99.999% purity) at 30 mL/min for 10 min. The volatiles were carried by the gas flow and trapped onto the Tenax TA adsorbent. To minimize thermal degradation and potential analyte loss, the entire purge-and-trap assembly was maintained at 4 °C, ensuring rapid cooling of outflowing gases from the glass vial. Post-collection, the Tenax TA liner was directly transferred to a gas chromatography–mass spectrometry (GC-MS) system (Agilent) for thermal desorption and analysis (See Figure S1B).

### GC-MS Analysis

Volatile compounds were analyzed using GC-MS equipped with a DB-17ms capillary column (30 m × 0.32 mm). The inlet temperature was set to 280 °C. The GC temperature program consisted of an initial hold at 40 °C for 2 minutes, followed by a temperature ramp from 40 °C to 130 °C at 5 °C/min (Segment 1), then from 130 °C to 250 °C at 20 °C/min (Segment 2), with a final hold at 250 °C for 3 minutes (Segment 3). Helium was used as the carrier gas at a constant column flow rate of 1 mL/min. MS acquisition was performed using the standard spectra tuning mode. Mass spectral data were analyzed using the National Institute of Standards and Technology (NIST) library.

### Sample preparation for chemotaxis assays

Worm-conditioned media, the native VSPs from *C. elegans* hermaphrodites and *C. remanei* females in M9 buffer were prepared and used for the chemotaxis assay according to the methods of Leighton *et al*. [12] and Wan *et al* [28]. For the initial chemical screens (Figures 1 and 2), test compounds were prepared at 1,000-fold and 10,000-fold dilutions. Both the 1,000-fold and 10,000-fold dilutions were prepared to ensure that the final ethanol concentration remained constant at 0.01%. Control solutions were prepared using the same ethanol and M9 buffer ratios, but without the test compound.

**Figure 1:**
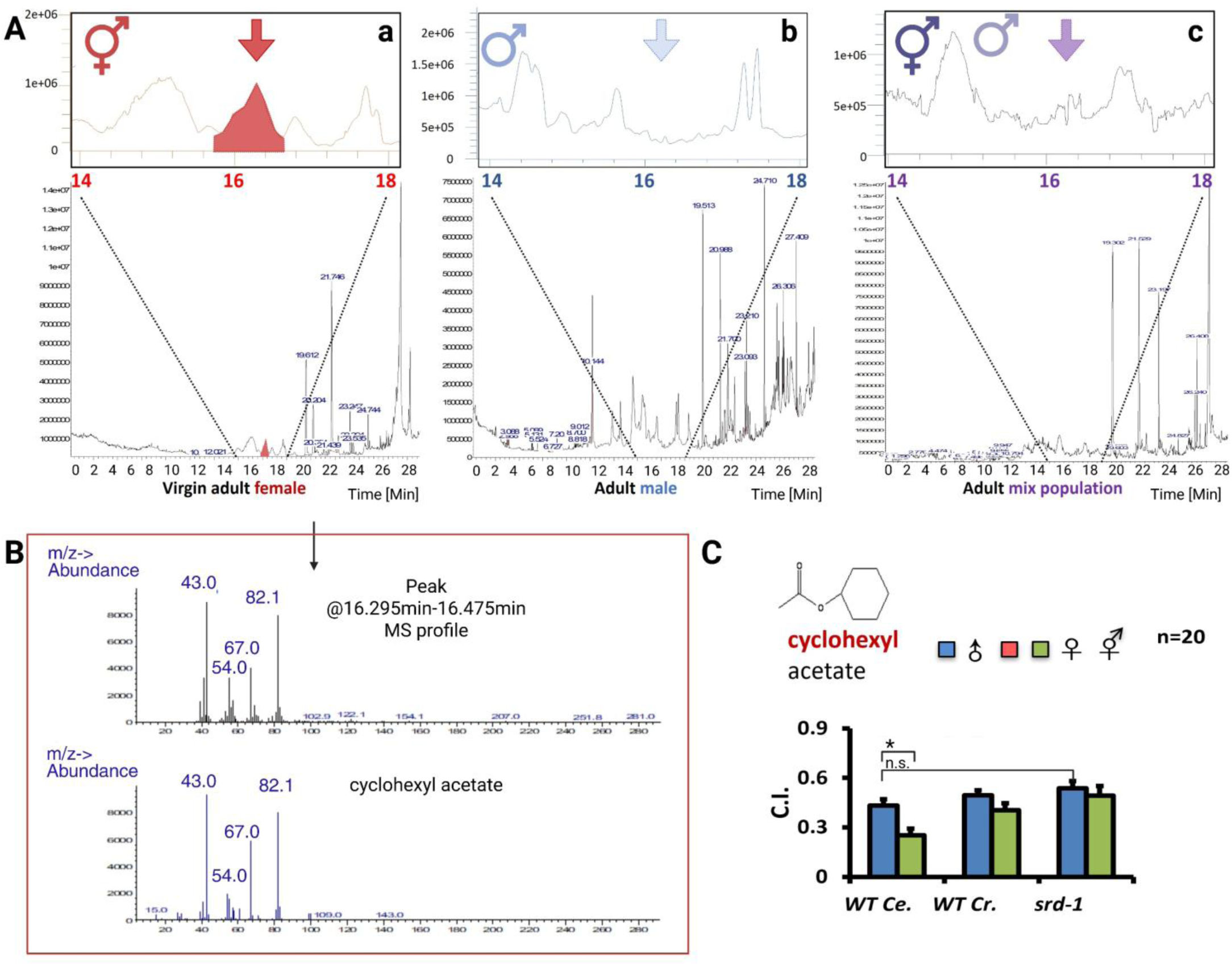
(A) Gas chromatography (GC) profiles of extracts from *C. remanei* (EM464) adult worms. (a) Virgin female extract showing a specific peak (red arrow) at retention time 16.295–16.475 min. (b) Male extract lacking the female-specific peak. (c) Mixed-population extract confirming the absence of the peak in the 16.295–16.475 min window. Mass spectrometry (MS) analysis of this region in (b) and (c) showed no match to the female-specific compound. (B) Mass spectrometry (MS) validation. Upper panel: MS profile of the female-specific peak (16.295–16.475 min). Lower panel: Reference MS profile of cyclohexyl acetate standard. All major peaks align. (C) Chemotaxis indices of wild-type *C. elegans* and *C. remanei* in response to 1000-fold diluted cyclohexyl acetate. Error bars denote standard error. * Indicates a significant difference (P < 0.05).

**Figure 2.**
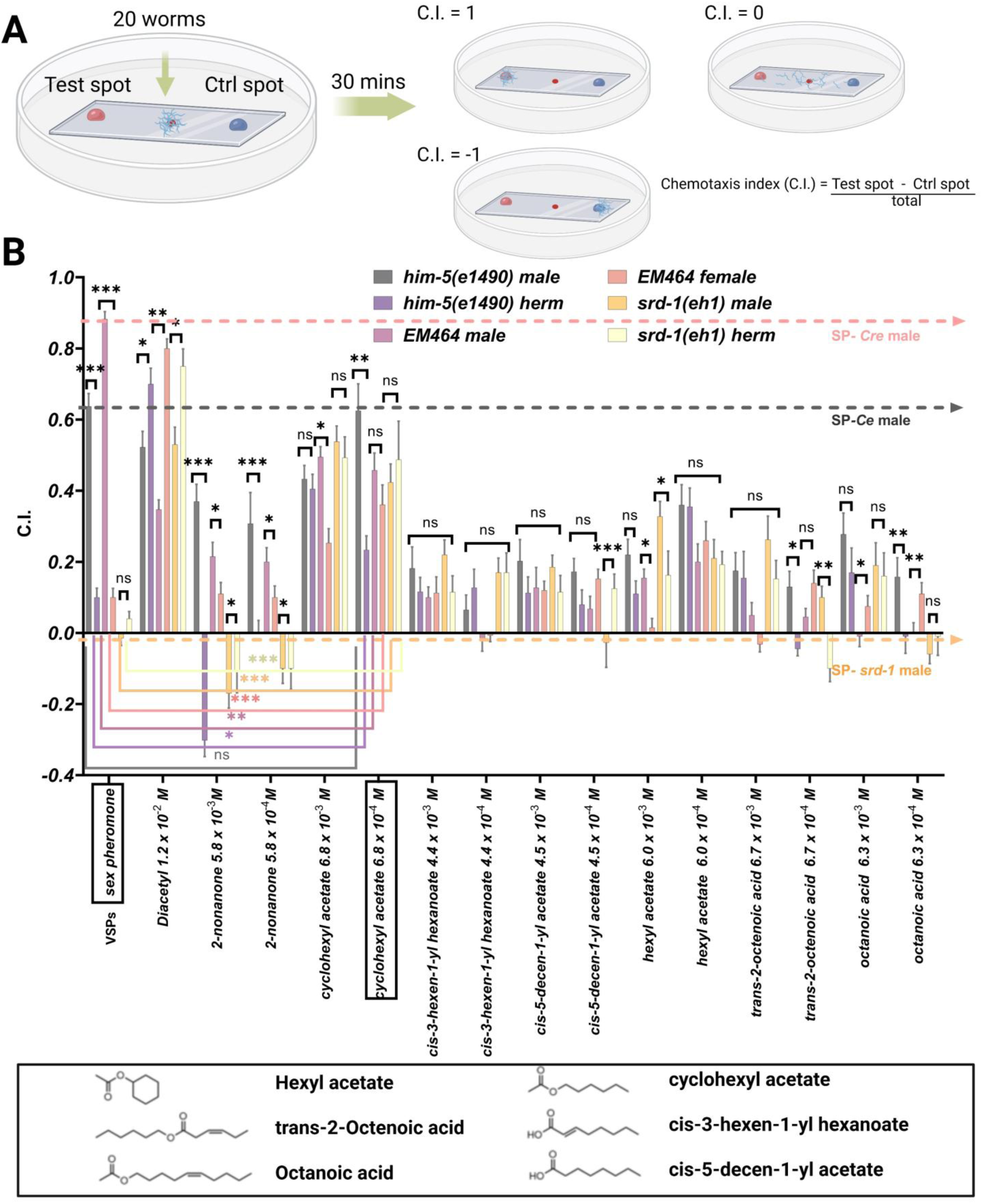
Chemotaxis index of wild-type *C. elegans* and *C. remanei* to VPS and volatile organic compounds at 1:1,000 and 1:10,000 dilutions. Chemotaxis indices (C.I.) of *him-5(e1490) C. elegans,* EM464 *C. remanei* and *srd-1* mutant strain of both sexes in response to different volatile compounds. The graph shows the chemotaxis indices of the test strains in response to VPS and various chemical compounds at different concentrations (1:1,000 and 1:10,000), including cyclohexyl acetate, diacetyl acetate, cis-3-hexen-1-yl hexanoate, cis-5-decen-1-yl acetate, hexyl acetate, 2-octenoic acid and octanoic acid. Positive indices indicate attraction, whereas negative indices indicate repulsion. Diacetyl and 2-nonanone served as the attractant and repellent positive controls, respectively. Sex pheromone (VSPs) served as the positive control. Chemotaxis index (C.I.) = (number of nematodes in the test spot) - (number of nematodes in the control spot) /total number of worms. The error bars represent the standard error. *, **,*** indicate a significant difference (*P <* 0.05, 0.005 and 0.0001, respectively).

For the concentration-response, mutant assays and adaptation assay (Figures 3-5 and S1), a serial dilution series of cyclohexyl acetate (CA) and 2-cyclohexen-1-ol (2CH) was prepared, starting from a 50% (v/v) stock solution in ethanol. Serial dilutions were made in M9 buffer to generate seven final concentrations ranging from 3.4 × 10⁻² M to 3.4 × 10⁻⁸ M.

**Figure 3.**
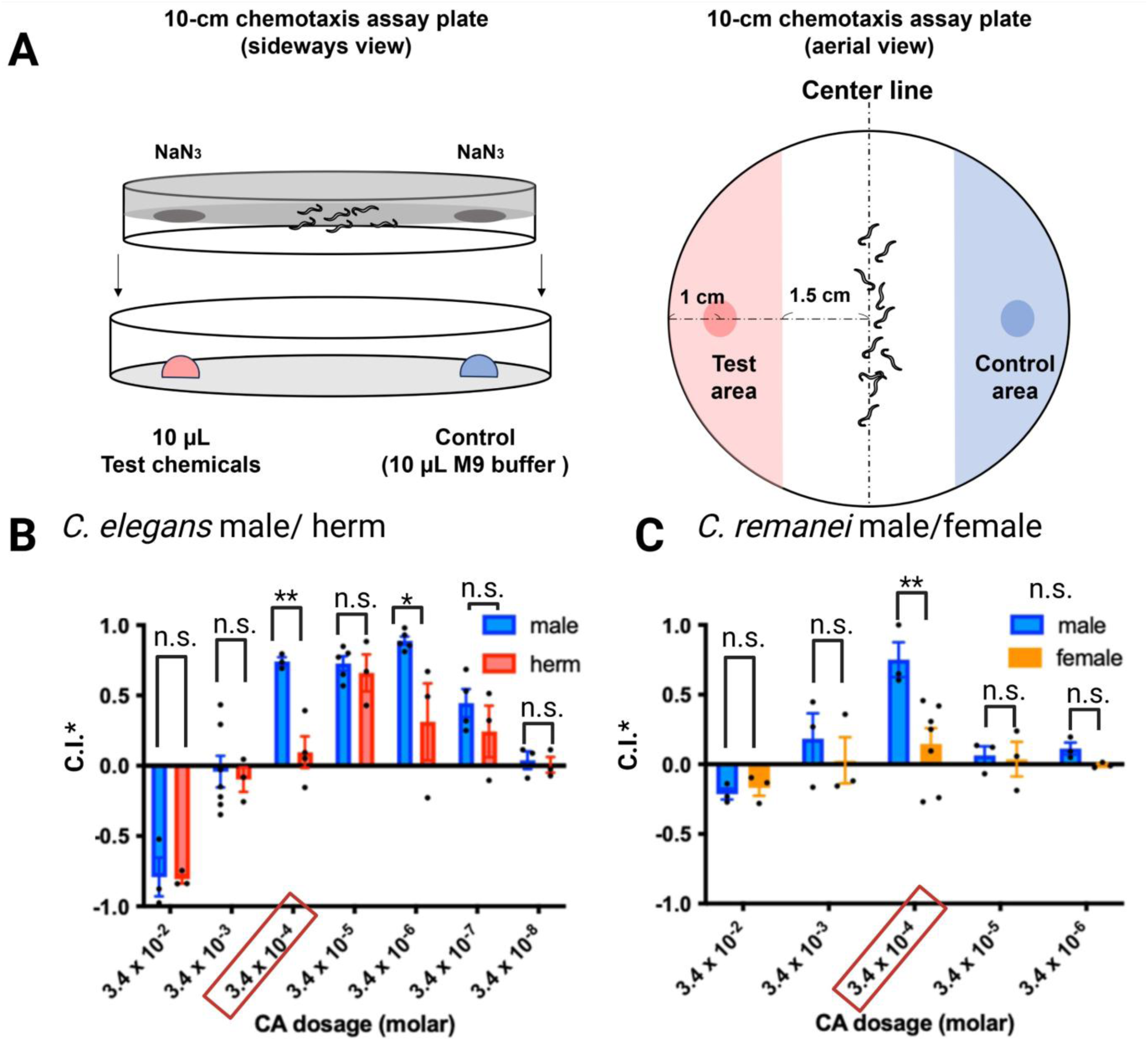
Sex-specific and concentration-dependent responses to cyclohexyl acetate in *C. elegans* and *C. remanei*. (A) Schematic diagram of the 10-cm plate chemotaxis assay. (B) Chemotaxis responses of *C. elegans* males and hermaphrodites across a dilution series of CA from 3.4 × 10⁻² M to 3.4 × 10⁻⁸ M. (C) Chemotaxis responses of *C. remanei* males and females across 3.4 × 10⁻² M to 3.4 × 10⁻^6^ M range. Chemotaxis index* (C.I.*) = (number of nematodes in the CA zone) - (number of nematodes in the control zone)/{(number of nematodes in the CA zone) + (number of nematodes in the control zone)}. The error bars indicate the standard error. *, ** indicate a significant difference in the t-test (*P <* 0.05 and 0.005, respectively)

**Figure 4.**
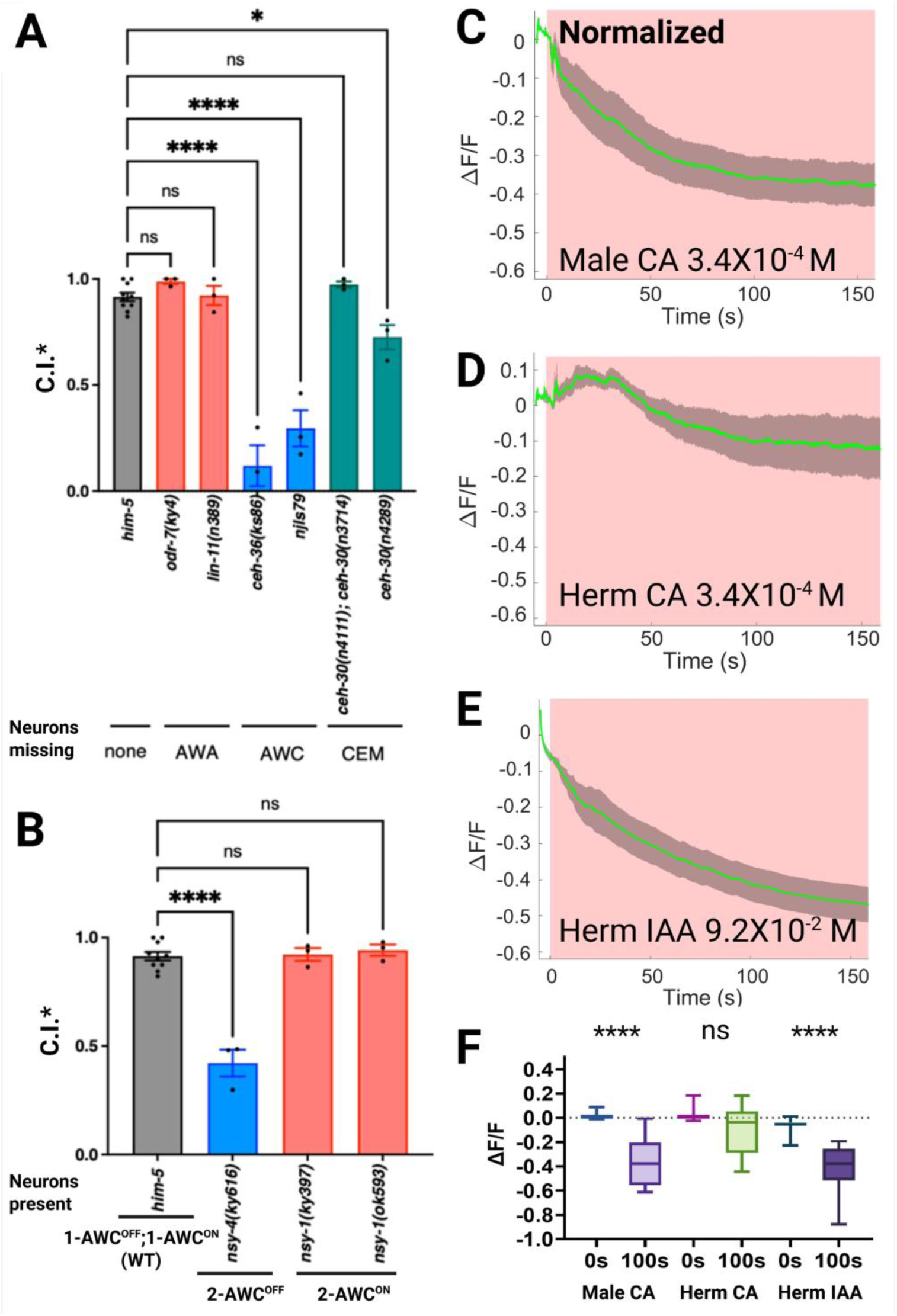
Chemotaxis deficits in *C. elegans* pheromone-detecting neuron mutants and sex-dimorphic AWC_on_ neuronal activity in response to cyclohexyl acetate. (A) Male attraction of *C. elegans* AWA, AWC, and CEM neuron-defective mutants to CA at 3.4 ×10^-6^ M. (B) Male attraction of *C. elegans* 2-AWC^on^ and 2-AWC^OFF^ mutants to CA at 3.4 ×10^-6^ M. Error bars indicate standard error. *, **** indicate a significant difference (*P <* 0.05 and 0.0001, respectively) in C.I.* between the mutants and *him-5* control based on Dunnett’s test. (C to E) Activity of AWC^on^ neurons in *C. elegans* adult males or hermaphrodites (herm) in response to CA and IAA. GCaMP6s signals were normalized to coelomocyte GFP to correct for eliminated photobleaching effect, which was measured via the same setup. Green line: ΔF/F average. Gray shading: s.e.m. envelope. The red bar indicates the stimulation time. n = 14 and 9 for the CA test in males and hermaphrodites, respectively, and n=15 for the IAA-positive control. (F) GCaMP6s signal intensity of *C. elegans* males and hermaphrodites subjected to CA stimulation and hermaphrodites subjected to IAA stimulation at 0 s and 100 s after stimulus delivery.

**Figure 5.**
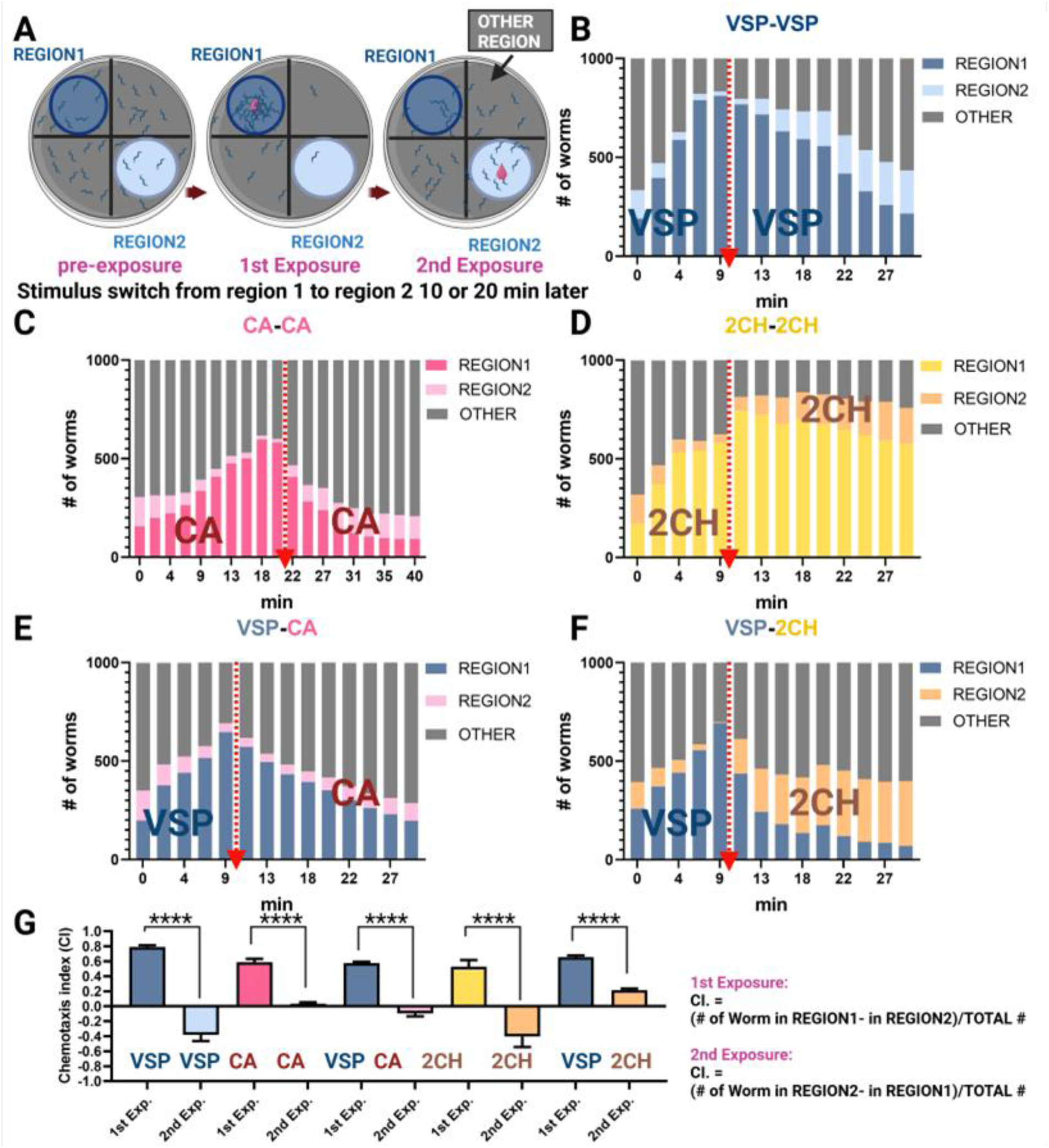
Adaptation assay schematic and stimulus-specific attenuation of male chemotaxis in *C. elegans* to VSP media and volatile compounds with cross-adaptation effects. (A) A schematic overview of the adaptation assay procedure. One-day-old adult *him-5* males were transferred to assay plates and were allowed to distribute for 10 minutes. Sequential stimuli (VSP, CA, or 2CH) were applied to designated regions. First, apply the stimulus to Region 1. Then, remove the stimulus from Region 1 and apply the second stimulus to Region 2. WormLab software recorded and quantified worm positions at 2.22-min intervals. (B) (C) Males pre-exposed to worm-conditioned VSP media or CA are no longer attracted by the same concentration of VSP or CA, respectively. (D) Males pre-exposed to 2CH are not attracted by the same concentration of 2CH in Region 2. (E) CA in the second exposure test no longer attracted males that were pre-exposed to VSP. (F) 2CH still attracted males that were pre-exposed to SP. (G) Chemoattraction index was measured after the first and second exposures. Significance was determined by a paired t-test: ****P<0.0001. The error bars indicate the SDs. The CA concentration is 3.4 × 10-^4^ M, and the 2CH concentration is 1.4 × 10^-3^ M. The dashed line with an arrowhead indicates the chemical switch.

### Chemotaxis assays

Chemotaxis assays were performed according to Leighton et al. [29] and Wan et al [19, 30] with slight modifications. Both the health and age of the worms affect their behavioral response to pheromones. The worms used in the chemoattraction assay were first bleached and synchronized to ensure that they were clean and at the same developmental stage. Worms were separated from individuals of the opposite sex by picking during the L4 stage and were then fed for another day. The worms were then rinsed from the plate with M9 and then washed twice with M9 buffer and then twice with ddH2O. The chemotaxis assay was carried out at room temperature in a 6- or 10-cm Petri dish prepared with chemotaxis media (2.0% agar, 1 mM MgSO4, 1 mM CaCl2, and 25 mM KH2PO4 (pH 6.0))

For the chemical screen (Figures 1 and 2), we adapted a 6-cm plate assay system. A glass coverslip pre-coated with chemotaxis media agar was placed in a Petri dish. The experimental site added 2 μL of test compound (e.g., VSPs) and 2 μL of 1 M sodium azide on the agar surface. Sodium azide was used to immobilize worms upon arrival. The control site added 2 μL of control buffer and 2 μL of 1 M sodium azide on the plate. For 6-cm Petri dishes, the standard distance between the center and each test site was set to 1.5 cm. To initiate the assay, 20 healthy, actively moving worms were manually picked and released simultaneously at the center point. Immediately afterward, 2 μL of the test compound and control solution were added to the corresponding spots. The plate was gently closed and placed undisturbed in a room temperature-stable environment. After 30 minutes, the number of worms at each site was counted. Worm picking was completed within 1–2 minutes, and the entire setup process was finished within 5 minutes to ensure assay consistency.

For chemical concentration-response and mutant assays (Figures 3 and 4), a modified lid-based chemotaxis protocol was used. Approximately 100 worms were pipetted directly onto the center of each chemotaxis plate. Two 1 μL drops of 1 M sodium azide were applied to opposite sides of the agar surface. A 100 μL aliquot of cyclohexyl acetate (CA) at concentrations ranging from 3.4 × 10⁻² M to 3.4 × 10⁻⁸ M was placed on the inner lid above sodium azide spot. For VSP assays, 10 μL of VSP extract was applied to the lid above the sodium azide site, with 10 μL of M9 buffer used as a control above the opposite sodium azide site (Figure S1). Plates were incubated for 30 minutes before scoring, and chemotaxis indices were calculated based on worm distribution. The assay plate was then placed in a small box to eliminate the influence of light. The assay was run for at least 1 hr. If >20% of the nematodes were still alive after 1 hr, we extended the incubation time until >80% of the nematodes tested died, as determined by periodic checking. This took between 1 hr. and 4 hr. All chemotaxis assays were repeated at least three times.

### Calcium imaging

Intracellular calcium signals were measured by monitoring Ca^2+^ binding to a calcium indicator, GCaMP6s; this binding increases the intensity of green fluorescence in cells. Calcium imaging was performed in a disposable worm-mounting and chemical-loading device, the thin-layer microdiffusion (TLMD) unit[18]. For the neural-excitation assays, we used a Nikon STORM spinning-disc microscopy system, which comprised a Nikon Ti2-E motorized inverted microscope, a Yokogawa CSU-X1 spinning disk with microlenses, and two EMCCD cameras for detection. A Nikon 40X dry objective lens was used. Time-lapse images were acquired at ∼8 frames/s for 150 sec at a 488 Hz laser intensity of 20%. The test samples were added manually at ∼ 5 s after filming started. Data analysis and video stacking were performed in ImageJ. Both AWC-neuron cell-body and dendrite regions near the cell-body were selected and analyzed. Coelomocyte GFP: Serves as an internal control for photobleaching correction. The normalized GCaMP6s signal intensity was calculated by normalizing the recorded intensity at each time point against the average coelomocyte GFP fluorescence intensity photobleach rate measured by the same setup. The fluorescence changes were normalized to the pre-stimulation background (before time 0), and the relative intensity values ΔF/F are presented here. The data is presented as the means ± s.e.m. All the data were analyzed with GraphPad Prism 6 and Microsoft Excel. Figures were plotted via custom MATLAB code [29]. Green line: ΔF/F average. Gray shading: s.e.m. envelope. The red or yellow bar indicates the stimulation time.

### Adaptation assay in *C. elegans*

Before the start of the assay, all of the one-day-old adult worms of CB4088*: him-5(e1490)* (standard male-rich reference strain) to be tested were washed from the OP50 seeded plate with M9 buffer and transferred to a clean plate without food. This was to ensure that all bacteria were removed from the body surface of the worms so that the worms would not be affected by the attraction of food during the assay. Before the test, the worms were examined via a chemoattraction assay with a positive control: 1.2 × 10^-2^ M (1:1000) diacetyl (a well-documented *C. elegans* attractant diluted in 10% ethanol) and VSP extracted from *C. elegans fog-2* virgin adult females. Only the worms that passed this quality test were used for further examination. A 9-cm Petri dish was prepared with 48 mL of adaptation assay agar (1.5% agar, 25 mM NaCl, 1.5 mM Tris-base, and 3.5 mM Tris-Cl). A circle of radius 1 inch was marked in the center of the Petri dish and divided into four quadrants, with two stimulus regions delineated at the centers of two opposite quadrants (labeled regions 1 and 2). All the markings were marked on the bottom of the Petri dish. A group of 100 synchronized and M9 precleaned worms were placed in the center starting site of the Petri dish and allowed to be distributed on the plate over a 10-minute period.

In the first stimulus exposure step, a 2 µl aliquot of VSP from *C. elegans fog-2* females, 3.4 × 10^-4^ M (1:20000) cyclohexyl acetate (CA) or 1.4 × 10^-3^ M (1:5000) 2-cyclohexylethanol (2CH) solution, was placed on the lid corresponding to the center point of region 1. The lid was closed, and the assay was recorded via WormLab (MBF Bioscience). At the 10- or 20-minute mark, the stimuli on the lid of Region 1 were removed, and the second exposure step was initiated by placing the second stimulus sample on the lid of the center point of Region 2. The recording continued for 20 minutes. WormLab software was used to perform data analysis, with the number of worms located within regions 1 and 2 counted every 1000 frames equating to approximately 2.22 minutes, where the frame rate was 7.5 fps. We administered the first exposure solution and conducted recordings for either 10 or 20 minutes, corresponding to either 4500 or 9000 frames. This resulted in 5 and 10 data points (including T0), respectively. Subsequently, we temporarily paused the recording for several seconds to switch the exposure solution. Upon restarting the recording following the introduction of the second exposure solution, we continued the recording for an additional 20 or 30 minutes, equivalent to 9000 or 13500 frames, thereby yielding 9 and 14 data points, respectively. In the VSPs adaptation assay (Figure S3B), the duration of the second exposure was prolonged to 30 minutes. This additional time was vital for determining the point at which no further significant changes in the chemoattraction index were noted, guiding the timing for subsequent experiments.

The chemotaxis index (C.I.) was calculated as the difference between the number of worms in the experimental region and the control region divided by the total number of tested worms. A C.I. value of 1.0 indicates that all the worms are attracted to the experimental spot, and a C.I. value near zero means that the test material is not attractive to the worms. Assays were performed on three separate days with freshly prepared samples. The sample size was 10 assays (1000 animals) for each test.

## Result

### Identification of a female-specific volatile compound in *Caenorhabditis* via GC-MS

Gas chromatography (GC) analysis of one-day-old adult *C. remanei* identified a female-specific compound in virgin female extracts, marked by a distinct peak at a retention time of 16.295–16.475 minutes (Figure 1A, panel a). This peak was absent in both males (Figure 1A, panel b) and mixed-population extracts (Figure 1A, panel c), indicating sex- and reproductive-state specificity. Mass spectrometry (MS) analysis of the corresponding GC region in male and mixed samples confirmed the absence of this compound. Virgin females of *Caenorhabditis* species (*C. elegans, C. remanei, C. briggsae and C. brenneri*) secrete VSPs that attract conspecific and heterospecific males [15]. Prior studies revealed that VSP production in *C. remanei* exceeds that of *C. elegans* by at least 20-fold [12, 15, 28]. Given the GC-MS requirement for large-scale sample preparation (5,000 virgin females *C. remanei* to achieve detectable signal-to-noise ratios), *C. remanei* was selected for VSPs extraction. Mass spectrometry profiling of this female-specific peak yielded a fragmentation pattern most closely matching that of the cyclohexyl acetate (CA) standard (Figure 1B). While this does not definitively confirm the identity or structural similarity of the compound, CA received the highest match score among all tested chemical standards in the NIST library.

### Cyclohexyl acetate elicits male-specific attraction

We used the chemoattraction assays adopted from Wan *et al*. [18, 28] with minor modifications. Chemoattraction assays using 1000-fold diluted CA demonstrated significant attraction in both sexes of *C. elegans* and *C. remanei*, with robust positive chemotaxis index (Figure 1C). However, the known VSPs chemoreceptor *srd-1* mutant males did not show a significant reduction in chemotaxis toward cyclohexyl acetate, suggesting that detection of this compound does not depend on the SRD-1 chemoreceptor. Sex pheromone blends within a species often consist of mixtures of structurally related compounds [5, 30-34]. For example, the cabbage looper produces a blend in which the major component is (Z)-7-dodecenyl acetate, accompanied by several minor acetate esters [30]. This phenomenon prompted us to screen chemicals that are structural variants of cyclohexyl acetate.

Then we evaluated a series of chemoattraction assays using a set of six compounds structurally related to cyclohexyl acetate but varying in hydrocarbon chain length, unsaturation, and the presence or absence of the acetate function group. Both species exhibit similar responses to most odors tested, with the exception of the repellent control 2-nonanone, which elicited a repellent response only in *C. elegans* males. Additionally, cis-3-hexen-1-yl hexanoate, hexyl acetate, trans-2-octenoic acid, and octanoic acid at specific concentrations induced slight attraction in *C. elegans* but not in *C. remanei*. Among those chemicals, CA functions as a strong concentration dependent male-specific attractant (Figure 2). EM464 *C. remanei* and CB4088: *him-5(e1490) C. elegans* were used as a standard male-rich reference strain. For *him-5(e1490)* males, CA at a concentration of 6.8 × 10^-3^ M resulted in chemotaxis index (C.I.) of 0.433 ± 0.038. The response increased at a lower concentration of 6.8 × 10^-4^ M, with a C.I. of 0.625 ± 0.075. This response was comparable to that of the positive control (VSPs, C.I. = 0.637 ± 0.036). In contrast, *him-5(e1490)* hermaphrodites also showed attraction to CA at higher concentrations, with a C.I. of 0.405 ± 0.041 at a concentration of 6.8 × 10^-3^ M, and a decreased response, with a C.I. of 0.233 ± 0.039 at 6.8 × 10^-4^ M. These results confirm that CA is a potent male-specific attractant at certain concentration. We evaluated the role of SRD-1, a key receptor for VSP [18], via *srd-1(eh1)* mutants (Figures 1 and 2).

However, the robust response observed in *srd-1* mutants of both sexes, which is equivalent to *him-5* response, suggested that CA detection involves *srd-1*-independent pathways integrated with other neural mechanisms. These data suggest that the unknown compound may share certain fragmentation features with CA, and we therefore interpret CA as a putative structural analog or mimic of a naturally occurring volatile component. Additionally, the observation that CA elicits robust male attraction in VSPs’ major chemoreceptor *srd-1* null mutants, suggests that CA is not a VSPs major component, it acts as a structural mimic of natural VSP components and functions via an alternative pathway.

### Cyclohexyl acetate is a concentration-dependent male attractant

To investigate the functional relevance of cyclohexyl acetate (CA) as a candidate mimic of female VSPs, we performed quantitative chemotaxis assays across a broad concentration range, using a protocol adapted from Leighton et al. [12] with minor modifications (see Methods and Figure 3A). CA was tested at concentrations spanning seven orders of magnitude (3.4 × 10⁻² M to 3.4 × 10⁻⁸ M) to characterize dose-dependent responses in adult *C. elegans* and *C. remanei*.

In *C. elegans*, both males and hermaphrodites displayed concentration-dependent responses to CA, but with distinct behavioral profiles. At high concentrations (3.4 × 10⁻² M), both sexes showed aversion, suggesting that CA is aversive above a threshold. At intermediate concentrations (3.4 × 10⁻⁴ M to 3.4 × 10⁻⁷ M), males exhibited robust attraction, with a chemotaxis index* (C.I.*) peaking near 0.6 (Figure 3B). Hermaphrodites showed a significantly weaker response at 3.4 × 10⁻⁴ M and 3.4 × 10^⁻6^ M. *C. remanei* males also showed significant attraction to CA, with peak responses at 3.4 × 10⁻⁴ M, whereas *C. remanei* females were not attracted by CA at any concentration tested (Figure 3C). Together, these findings demonstrate that CA elicits a sex-specific, species-specific, concentration-dependent response, with attraction occurring within an intermediate range and aversion emerging at higher doses.

Although the mechanism underlying aversion to higher concentrations of CA remains unclear, similar concentration-dependent aversions have been well documented for chemicals such as diacetyl, benzaldehyde, and isoamyl alcohol, which are also detected by AWA and AWC neurons [35-37]. At lower concentrations, AWA and AWC neurons trigger attraction, but at higher concentrations, ASH neurons mediate avoidance [35-37].

### Cyclohexyl acetate is perceived by *C. elegans* males via AWC^on^ neurons

Having established that CA is attractive to *C. elegans* and *C. remanei* males, we carried out chemotaxis assays with specific *C. elegans* mutants to identify the neurons responsible for CA perception. Both male-specific neurons (CEMs) and amphid sensory neurons (AWAs) have been implicated in VSP perception in *C. elegans* [15, 17-19]. Accordingly, we assayed males with mutations that affect the function of these neurons as well as that of an additional amphid neuron (AWC) responsible for volatile attractants [5, 17, 38]. Males of the *ceh-36(ks86)* and *njls79 [ceh-36p::cz::casp3, ceh-36p::casp3::nz, ges-1p::GFP](X)* strains, which lack functional AWC neurons [39, 40], were almost completely defective in their response to CA at 3.4 ×10^-6^ M (*p < 0.0001*, Dunnett’s multiple comparison test), whereas males of genotypes *odr-7(ky4)* and *lin-11(n389)*, which lack functional AWA neurons [41, 42], were attracted to the CA at the same level as wild-type (WT) males (Figure 3A). Males of *ceh-30(n4289)*, which lack functional CEM neurons [43], were slightly less attracted (*p < 0.05*, Dunnett’s multiple comparison test), suggesting that AWC neurons are primarily responsible for CA perception, with a possible contribution from CEM neurons. The two AWC neurons of *C. elegans*, designated AWC^on^ and AWC^off^, display molecular and functional asymmetry, as they sense partly different odors [44]. To determine whether both or only one of the two types of AWC neurons is required for CA perception, we performed chemotaxis assays using mutant lines. Mutant *nsy-4(ky616)* males, which exhibit a two-AWC^off^ phenotype [45], exhibited significantly less attraction (*p < 0.0001*, Dunnett’s multiple comparison test), whereas *nsy-1(ky397)* and *nsy-1(ok593)* mutants, which have two AWC^on^ neurons [46, 47], were attracted to the CA at the same level as WT males having both AWC^on^ and AWC^off^ neurons (Figure 3B). Therefore, AWC^on^ neurons are primarily responsible for CA perception.

To visualize the response of *C. elegans* AWC^on^ neurons to CA, we used the calcium indicator GCaMP6s for real-time visualization of sensory neuron excitation by monitoring the influx of calcium ions upon exposure to the chemical signal. We generated a transgenic worm strain carrying AWC^on^-specific regulatory sequences (P*str-2* promoter) that directly express the GCaMP6s reporter gene in AWC^on^. Previous studies have demonstrated that the calcium level in AWC^on^ neurons decreases upon odor addition [48-51], and as shown in Figure 3C and Figure S2, the fluorescence intensity in male AWC^on^ neurons largely decreases upon CA stimulation. Specifically, the GCaMP6s signal intensity decreased in the AWC^on^-neuron cell body, dendrites, and axons. The male AWC^on^-neuron GFP intensity decreased immediately after CA stimulation and rapidly decreased within the observation window of 2.5 min. CA is thus sensed by male AWC^on^ neurons. In contrast, hermaphrodites exposed to CA showed a weaker change in AWC^on^ neuron fluorescence intensity; the amplitude was one-fourth that of the male response (Figure 3D). The signal change was not significant and was more variable than that in the male AWC^on^ neuron response (Figure 3F). As a control, we tested another AWC-specific odorant, isoamyl alcohol (IAA), and found that the hermaphrodites of the imaging strain can respond to AWC-specific odorants (Figure 3E). The sexual dimorphism noted in AWC^on^ neuron activity aligns with our previous observations on AWA neurons’ responses to VSP containing natural pheromones [18].

### Cyclohexyl acetate functions as a female/hermaphrodite volatile sex pheromone mimic in *C. elegans*

To support our hypothesis that CA mimics a functional component of the VSPs produced by *C. elegans* hermaphrodites, we conducted adaptation assays with *C. elegans* males to the VSP extract from *fog-2* females, which contains the native VSPs [12], and to CA specifically (Figure 5). If prior exposure to VSPs does not reduce the male response to CA, it would suggest that CA is not mimicking VSPs. Conversely, if VSPs exposure diminishes the response to CA, it would support the idea that CA functions as a mimic of VSPs. The observed reduction in chemotactic response to CA following VSPs exposure serves to evaluate whether CA functions as a structural and functional mimic of the natural VSPs.

Initial exposure to both the natural VSP extract and CA strongly attracted male subjects, thus verifying their attractiveness to males in this behavioral assay context (Figure 5). Pre-exposure to both samples for 10 minutes significantly reduced male responsiveness, indicating olfactory adaptation to both the pheromone and the same concentration of CA, as evidenced in Figures 5B, 5C, and 5G. These behavioral experiments revealed that variations in odor preferences arise from pheromone-specific adaptation, suggesting that pre-exposure results in the rewiring of pheromone perception neural circuits. Not all odorants induce adaptation; for example, males pre-exposed to diacetyl (2,3-butanedione, DA) remain attracted to the same concentration upon re-exposure (Figure S3C).

To further explore a possible functional role of CA or a CA analog as a VSPs mimic, we conducted an experiment where males were first exposed to the natural VSPs for 10 minutes and then assessed for their chemotactic behavior toward CA. The underlying hypothesis being that if a chemical induces adaptation following pre-exposure and if this chemical is a constituent of a mixture, then prior exposure to this mixture should result in a reduced chemotactic response to this chemical, signaling olfactory adaptation, a behavior that has been demonstrated in *C. elegans* [51]. The results supported this hypothesis: pre-exposure to either the natural VSPs or CA led to a suppression of the chemotactic response to CA (Figures 5C, 5E, and 5G). We observed that males typically need approximately 16–20 minutes to respond robustly to CA, in contrast to the peak responses to other tested stimuli, which occur within 6–10 minutes. Therefore, the duration of the initial exposure was increased by an extra 10 minutes (Figure 5C). Males pre-exposed to CA retained attraction to the full VSP blend (Fig. S3A). This indicates that CA may only specifically mimic only one component of the VSP mixture. Sensory adaptation to CA selectively affects responses to some structural analog, but not the entire VSP mixture.

As a structurally analogous control to CA, 2-cyclohexylethanol (2CH) attracts males via AWC neurons [50] but is absent from natural VSP extracts. 2CH shows completely distinct mass spectral signature fragments to VSPs. The fragmentation profile of 2CH standard is missing the diagnostic m/z 43 fragment (characteristic of CA and natural VSP components) and it also has additional fragments at m/z 41, 55, 68, 69, 81, and 110. These spectral signatures were never detected in VSPs GC peak, confirming 2CH’s absence from the natural pheromone blend. We also performed a serial dilution assay and found that 2CH was more attractive at 1.4 × 10⁻³ M than at 3.4 × 10⁻⁴ M, which is the effective concentration for CA (Figure S1A). Therefore, we selected × 10⁻³ M for the adaptation assay, as it elicited the most robust attraction response in males (Figure S1A). We also tested hermaphrodite responses to different concentrations of 2CH. Hermaphrodites were more attracted to higher concentrations, but exhibited a repulsive response at the lowest concentration tested (3.4 × 10⁻⁶ M). Upon two exposures, 2CH did not produce a typical adaptation pattern. Instead, males are highly attracted by 2CH and remain predominantly at the site of initial exposure, resulting in limited migration to the alternate region and a minimal increase in the ‘other’ zone. These dynamics indicate that 2CH induces a strong site-retention effect (Figure 5D). Pre-exposure to VSP partially reduced the chemotactic response to 2CH (Figure 5F), though this attenuation was much less pronounced than the near-complete suppression observed for CA.

## Discussion

### Cyclohexyl acetate mimics a nematode sex pheromone to drive odor-specific olfactory adaptation via AWC neurons

This study identifies CA as a potent mimic of the natural *C. elegans* VSP and demonstrates that pre-exposure to CA or the natural pheromone mixture induces odor-specific adaptation in males. This study also demonstrated that AWC neurons mediate odor-selective adaptation rather than generalized sensory desensitization. These results show the specificity of chemosensory adaptation, allowing males to dynamically adjust their responses based on recent odor exposure. For example, pre-exposure to certain chemicals, such as 2CH, causes males to remain at the original exposure site. In contrast, pre-exposure to VSPs or CA induces adaptation, leading to a diminished response to subsequent exposures at the same concentration. Other chemicals, such as diacetyl, do not produce either retention or adaptation effects (Figures 5 and S3).

Notably, the CA mimics only a subset of the effects of natural pheromones. Unlike the full VSP, the CA does not rely on AWA neurons or the SRD-1 receptor for detection [18], suggesting that it replicates a single component of the multi-odorant pheromone system but is itself not a functional component of the VSP [18]. This highlights the intricacy of the *C. elegans* chemical communication system, where multiple pheromone components likely act in synergy to elicit the full behavioral response [2, 34, 52]. The identification of CA raises questions about its biological origin: while its role as a pheromone mimic is clear, it remains unresolved whether CA is an endogenous product of *C. elegans*, a microbial metabolite, or an environmental compound. This ambiguity highlights the need to investigate the natural occurrence and ecological relevance of CA, particularly given its potential to act as a decoy signal in habitats where microbes or other nematodes might produce structurally similar volatiles.

CA triggered variable GCaMP6s signal excitation in some hermaphrodite AWA neurons, whereas, in males, it elicited a steadier and stronger GCaMP6s signal response. These results are similar to those we previously reported for VSPs [19]: VSPs elicited an unstable, receptor-dependent excitation of the GCaMP6s signal in hermaphrodite AWA neurons, whereas, in male AWA neurons, the VSP evoked a more stable receptor-dependent GCaMP6s signal with a larger amplitude.

### Potential Implications for Cross-Species Interactions

While our study demonstrates that CA elicits male-specific attraction in *C. elegans* and *C. remanei*, its broader ecological significance remains to be explored. The observed difference in behavioral responses between *C. elegans* hermaphrodites and *C. remanei* females may reflect underlying differences in their reproductive strategies. *C. remanei,* a dioecious species, is subject to stronger sexual selection pressures that likely favor the evolution of strict sex-specific signaling mechanisms. In contrast, *C. elegans* is androdioecious, and hermaphrodites rarely outcross, reducing the selective pressure for female-specific pheromone responses.

The structural resemblance of CA to known nematode signaling molecules raises questions about potential cross-species interactions, as observed in other systems. For instance, nematophagous fungi exploits nematode-derived ascarosides to trigger trap formation, and plant-parasitic nematodes use pheromonal cues to facilitate beetle-mediated dispersal [23, 25]. If CA or structurally similar compounds are present in natural environments, they could be co-opted by predators as decoy signals or by symbiotic microbes to modulate host behavior. However, these scenarios remain speculative and require validation in ecologically relevant contexts. Notably, the endogenous source of CA has yet to be identified. To address these gaps, future studies should investigate whether CA is produced by soil-dwelling microbes or nematodes, assess its effects on non-*Caenorhabditis* species, and characterize the chemical composition of natural nematode habitats. Such efforts will be essential to clarify the ecological roles and evolutionary implications of CA in nematode communication.

### Role of *srd-1* in mediating male attraction to volatile cues

The *srd-1(eh1)* males presented a slightly decreased attraction response to CA (*him-5(e1490)* males, C.I.* = 0.628 ± 0.075; *srd-1(eh1)* males, 0.423 ± 0.052; at 6.8 × 10^-4^ M). A similar level of reduction was observed in response to other compounds, such as octanoic aid (*him-5(e1490)* males, C.I.=0.278 ± 0.060*; srd-1(eh1)* males, C.I.* = 0.190± 0.063; at 6.8 × 10^-3^ M); cis-5-decen-1-yl-acetate (*him-5(e1490)* males, C.I.=0.173 ± 0.037*; srd-1(eh1)* males, C.I.* = -0.028 ± 0.069; at 6.8 × 10^-4^ M dilution); and cis-3-hexenyl hexanoate (*him-5(e1490)* males, C.I.=0.182 ± 0.060*; srd-1(eh1)* males, C.I.* = 0.022 ± 0.042; at 6.8 × 10-3 M), among others. The consistent slight reduction in the chemotaxis index observed in *srd-1(eh1)* mutants across multiple volatile candidate derivatives suggests that *srd-1* specifically affects the detection of VSP structurally similar components. This pattern aligns with previous studies showing that *srd-1* mutants retain normal chemotaxis responses to pheromone-unrelated AWA-sensed volatile attractants [18], indicating that *srd-1* mutants do not have general defects in chemotaxis or mobility. This finding suggests a specific role for *srd-1* in recognizing particular chemical structures.

Overall, CA provides a fascinating lens for studying the complexity of sex pheromone communication in nematodes. While CA mimics a component of the *C. elegans* female VSP, its possible biogenic sources and potential for cross-species exploitation suggest critical questions for future research. These findings lay the groundwork for unraveling the ecological and evolutionary mechanisms driving volatile signaling in nematodes and their interactions with other organisms in their environment.

## Acknowledgments

We thank Dr. Shunji Nakano for previewing the manuscript and providing the worm strains. We thank Joshua N. Muller for assistance with behavior experiments. We thank WormBase for providing organized knowledge of *C. elegans* and the *Caenorhabditis* Genetics Center for strains. Dr. Michael Milligan provided invaluable assistance with the GC x GC-TOFMS analysis.

## Author contributions

PWS, RS, FCS, XW, YT, MG, DL, and CMS conceptualized the study. The investigation and data collection were carried out by XW, YT, and RS. Funding for the research was acquired by PWS, RS and FCS. Project administration was managed by MG, PWS, and RS. Supervision for the research was provided by PWS, RS and KLC. The original draft of the manuscript was written by XW, MG, FCS, PWS, and RS. All authors contributed to the review and editing of the manuscript.

## Data and materials availability

All the data are available in the manuscript or the supplementary materials.

---

**Figure S1.**
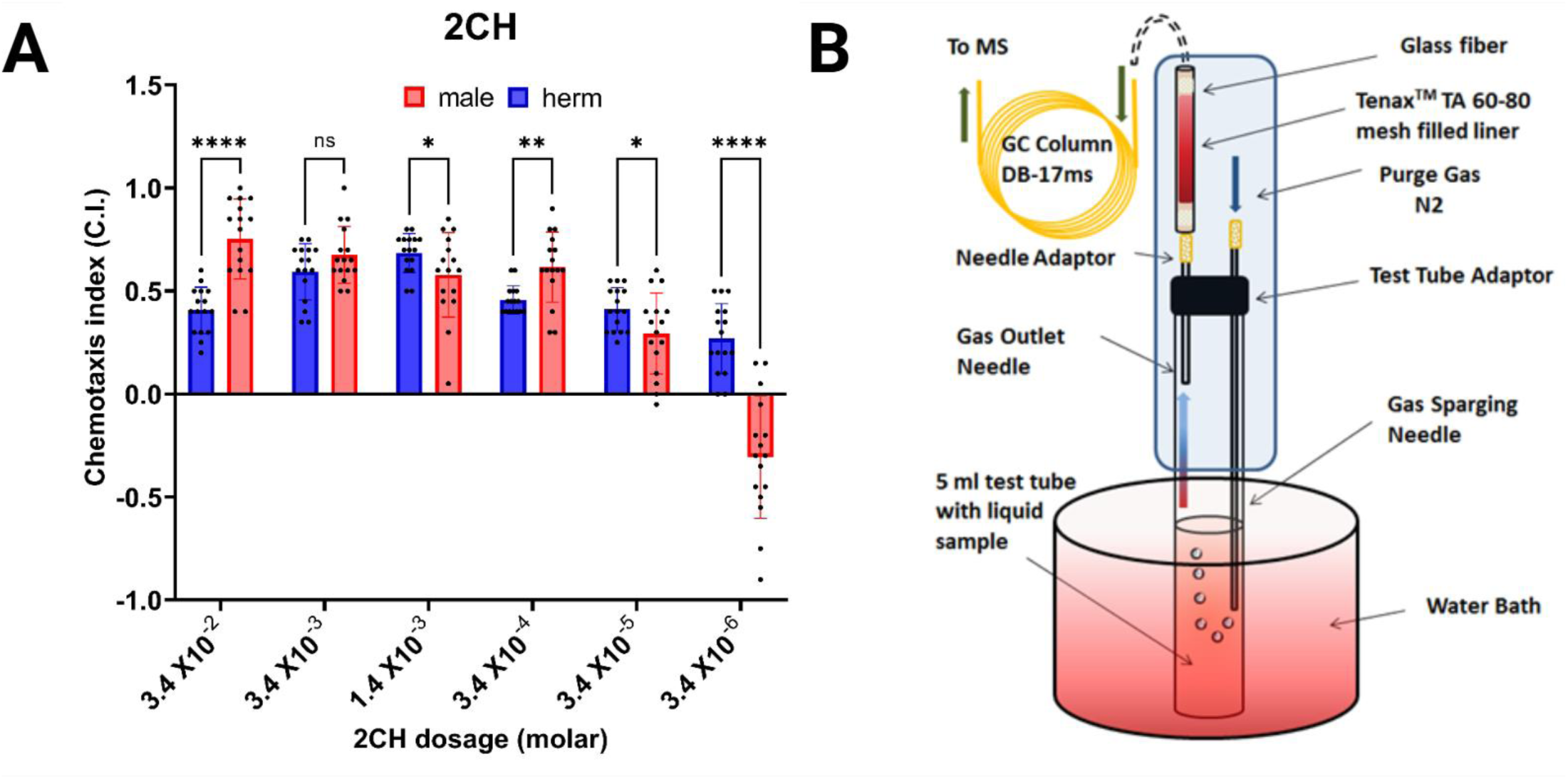
Chemotaxis assay design and response to 2CH in *C. elegans*. (A) Dose-dependent chemotaxis response of *C. elegans* males and hermaphrodites to different concentrations of 2CH. Error bars indicate standard error. *, **,**** indicate a significant difference (P < 0.05, 0.005, 0.000001, respectively). (B) Schematic of the GC-MS sample collection system. The trap and purge system is comprised of a liner filled with Tenax^TM^ TA (Poly [2, 6-diphenylphenylene oxide]) 60-80 mesh with a volatile component evaporation system. Pure and dry nitrogen gas passes through the collection system, and the volatile component is trapped in order to absorb material for further GC-MS analysis.

**Figure S2.**
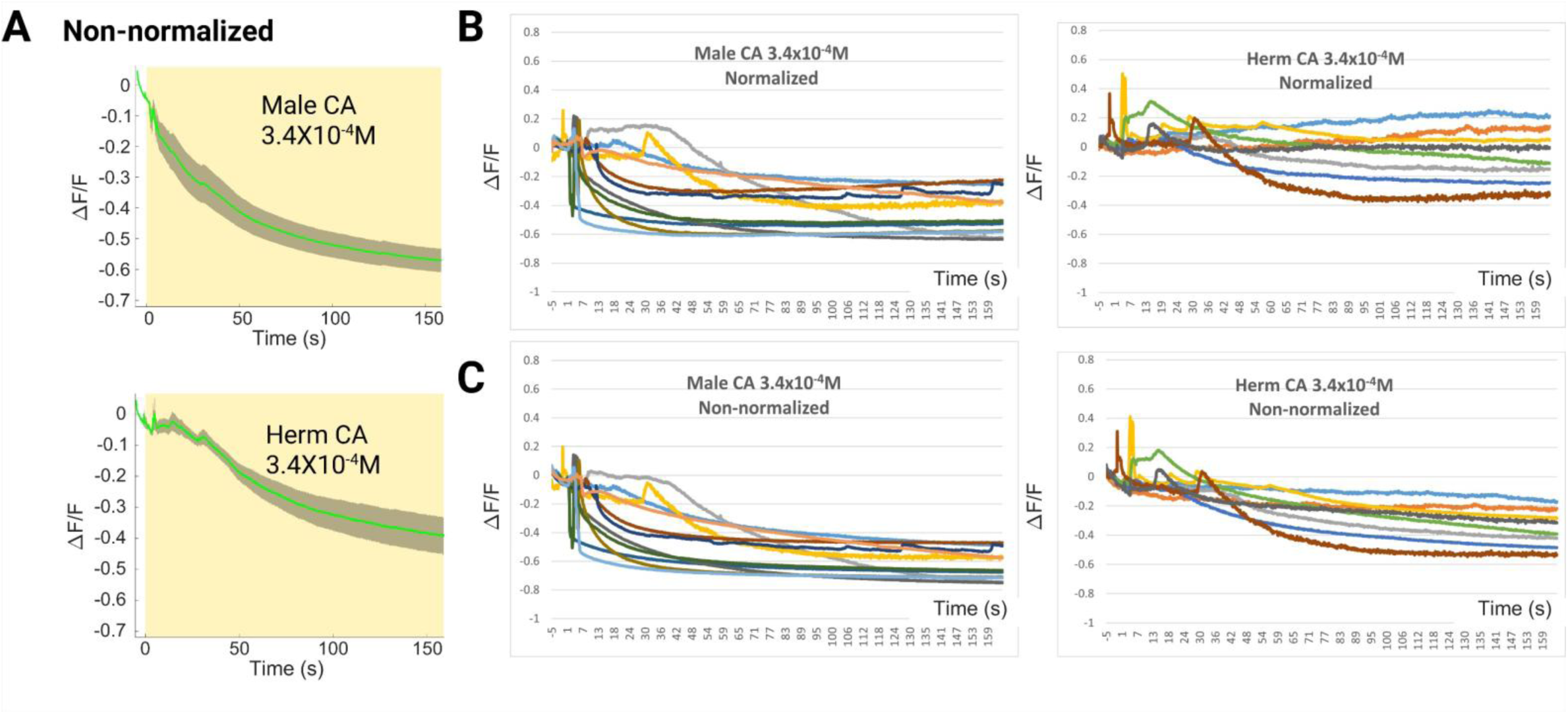
Cyclohexyl acetate activates a calcium response in male AWC^on^ neurons. Male AWC neurons are excited by the CA, and the response is much weaker in hermaphrodite AWC^on^ neurons. (A) Non-normalized data of GCaMP6s-induced GFP fluorescence was plotted without photobleaching rate normalization. Green line: ΔF/F_0_ average. Gray shading: s.e.m. envelope. The yellow bar indicates the stimulation time. (B) Normalized ΔF/F_0_ traces from individual animals. (C) Raw data: Non-normalized ΔF/F_0_ traces from individual animals.

**Figure S3.**
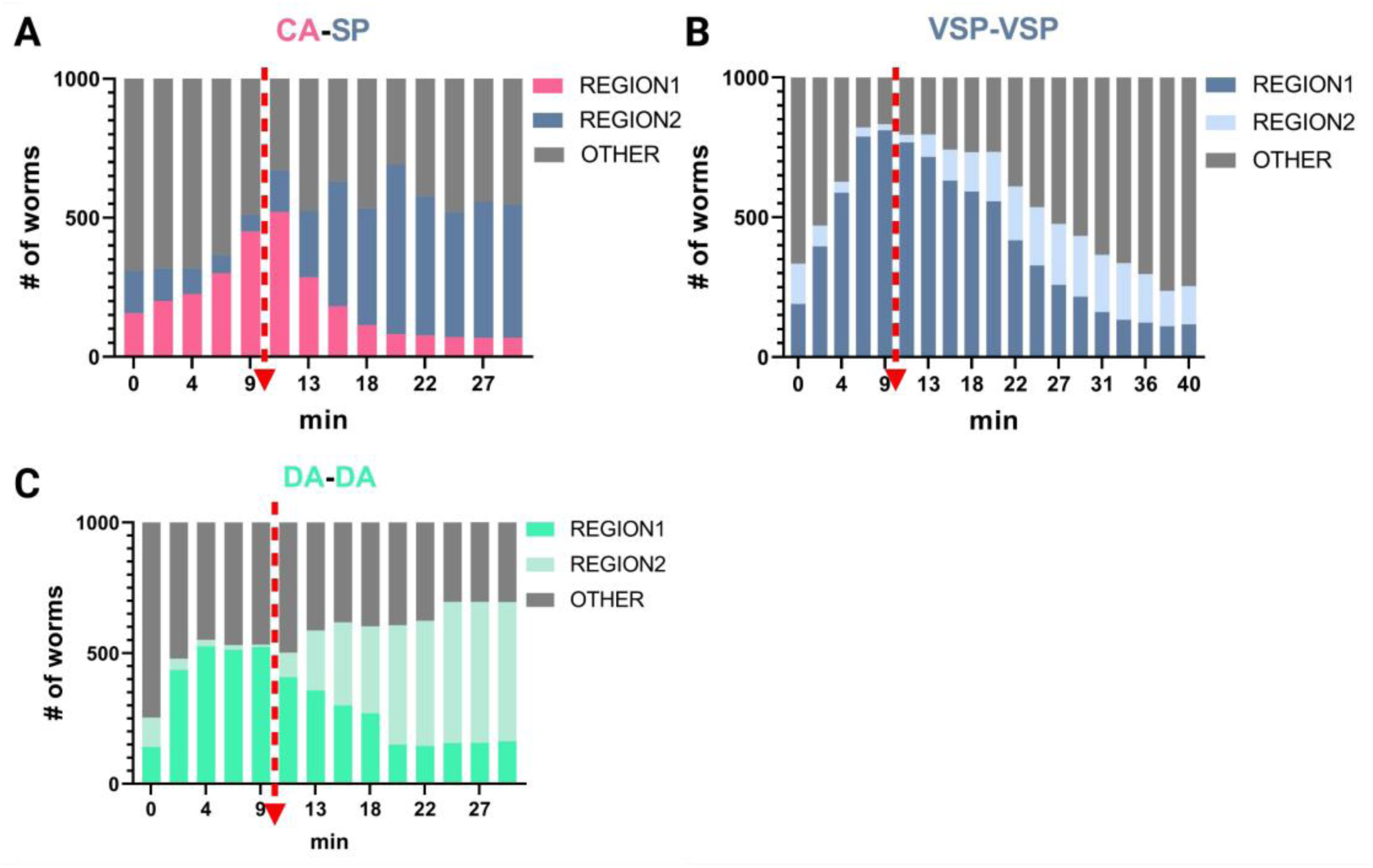
(A) Males pre-exposed to CA retained attraction to VSP. (B) Males pre-exposed to worm-conditioned VSP medium showed no attraction to the same VSP concentration. The chemoattraction index remained stable for 30 min, confirming adaptation persistence. (C) Males pre-exposed to diacetyl (2,3- butanedione, DA) remain attracted to the same concentration during subsequent exposure. The dashed line with the arrowhead indicates the timing of the chemical switch.

## References

1. Srinivasan J, Von Reuss SH, Bose N, Zaslaver A, Mahanti P, Ho MC, O’Doherty OG, Edison AS, Sternberg PW, Schroeder FC: A modular library of small molecule signals regulates social behaviors in Caenorhabditis elegans. PLoS biology 2012, 10(1):e1001237.

2. Srinivasan J, Kaplan F, Ajredini R, Zachariah C, Alborn HT, Teal PEA, Malik RU, Edison AS, Sternberg PW, Schroeder FC: A blend of small molecules regulates both mating and development in Caenorhabditis elegans. Nature 2008, 454(7208):1115-1118.

3. Golden JW, Riddle DL: A pheromone influences larval development in the nematode *Caenorhabditis elegans*. Science 1982, 218(4572):578-580.

4. Kaplan F, Srinivasan J, Mahanti P, Ajredini R, Durak O, Nimalendran R, Sternberg PW, Teal PE, Schroeder FC, Edison AS et al: Ascaroside expression in Caenorhabditis elegans is strongly dependent on diet and developmental stage. PLoS One 2011, 6(3):e17804.

5. Ludewig AH, Schroeder FC: Ascaroside signaling in C. elegans. WormBook: the online review of Celegans biology 2013:1–22.

6. Gallo M, Riddle DL: Effects of a *Caenorhabditis elegans* dauer pheromone ascaroside on physiology and signal transduction pathways. J Chem Ecol 2009, 35(2):272–279.

7. Baiga TJ, Guo H, Xing Y, O’Doherty GA, Dillin A, Austin MB, Noel JP, La Clair JJ: Metabolite induction of Caenorhabditis elegans dauer larvae arises via transport in the pharynx. ACS Chem Biol 2008, 3(5):294–304.

8. Wharam B, Weldon L, Viney M: Pheromone modulates two phenotypically plastic traits - adult reproduction and larval diapause - in the nematode *Caenorhabditis elegans*. BMC Evol Biol 2017, 17(1):197.

9. Golden JW, Riddle DL: A *Caenorhabditis elegans* dauer-inducing pheromone and an antagonistic component of the food supply. J Chem Ecol 1984, 10(8):1265–1280.

10. Park JY, Joo HJ, Park S, Paik YK: Ascaroside Pheromones: Chemical Biology and Pleiotropic Neuronal Functions. Int J Mol Sci 2019, 20(16).

11. Zhang X, Feng L, Chinta S, Singh P, Wang Y, Nunnery JK, Butcher RA: Acyl-CoA oxidase complexes control the chemical message produced by Caenorhabditis elegans. Proc Natl Acad Sci U S A 2015, 112(13):3955–3960.

12. Leighton DH, Choe A, Wu SY, Sternberg PW: Communication between oocytes and somatic cells regulates volatile pheromone production in Caenorhabditis elegans. Proceedings of the National Academy of Sciences of the United States of America 2014, 111(50):17905–17910.

13. Pungaliya C, Srinivasan J, Fox BW, Malik RU, Ludewig AH, Sternberg PW, Schroeder FC: A shortcut to identifying small molecule signals that regulate behavior and development in Caenorhabditis elegans. Proceedings of the National Academy of Sciences of the United States of America 2009, 106(19):7708–7713.

14. Choe A, von Reuss SH, Kogan D, Gasser RB, Platzer EG, Schroeder FC, Sternberg PW: Ascaroside signaling is widely conserved among nematodes. Curr Biol 2012, 22(9):772–780.

15. Chasnov JR, So WK, Chan CM, Chow KL: The species, sex, and stage specificity of a Caenorhabditis sex pheromone. Proceedings of the National Academy of Sciences of the United States of America 2007, 104(16):6730–6735.

16. Morsci NS, Haas LA, Barr MM: Sperm status regulates sexual attraction in *Caenorhabditis elegans*. Genetics 2011, 189(4):1341–1346.

17. White JQ, Nicholas TJ, Gritton J, Truong L, Davidson ER, Jorgensen EM: The sensory circuitry for sexual attraction in C. elegans males. Current Biology 2007, 17(21):1847–1857.

18. Wan X, Zhou Y, Chan CM, Yang H, Yeung C, Chow KL: SRD-1 in AWA neurons is the receptor for female volatile sex pheromones in *C. elegans* males. EMBO Rep 2019, 20(3).

19. Wan X, Zhou T, Susoy V, Park CF, Groaz A, Brady JF, Samuel AD, Sternberg PW: Efficient pheromone navigation via antagonistic detectors. bioRxiv 2024:2024.2011. 2022.624901.

20. Choe A, Stephan, Kogan D, Robin, Edward, Frank, Paul: Ascaroside signaling is widely conserved among nematodes. Current Biology 2012, 22(9):772–780.

21. Shinoda K, Choe A, Hirahara K, Kiuchi M, Kokubo K, Ichikawa T, Hoki JS, Suzuki AS, Bose N, Appleton JA et al: Nematode ascarosides attenuate mammalian type 2 inflammatory responses. Proc Natl Acad Sci U S A 2022, 119(9):e2108686119.

22. Manohar M, Tenjo-Castano F, Chen S, Zhang YK, Kumari A, Williamson VM, Wang X, Klessig DF, Schroeder FC: Plant metabolism of nematode pheromones mediates plant-nematode interactions. Nat Commun 2020, 11(1):208.

23. Hsueh YP, Mahanti P, Schroeder FC, Sternberg PW: Nematode-trapping fungi eavesdrop on nematode pheromones. Curr Biol 2013, 23(1):83–86.

24. Manosalva P, Manohar M, von Reuss SH, Chen S, Koch A, Kaplan F, Choe A, Micikas RJ, Wang X, Kogel KH et al: Conserved nematode signalling molecules elicit plant defenses and pathogen resistance. Nat Commun 2015, 6:7795.

25. Zhao L, Zhang X, Wei Y, Zhou J, Zhang W, Qin P, Chinta S, Kong X, Liu Y, Yu H et al: Ascarosides coordinate the dispersal of a plant-parasitic nematode with the metamorphosis of its vector beetle. Nat Commun 2016, 7:12341.

26. Artyukhin AB, Zhang YK, Akagi AE, Panda O, Sternberg PW, Schroeder FC: Metabolomic “dark matter" dependent on peroxisomal beta-oxidation in Caenorhabditis elegans. J Am Chem Soc 2018, 140(8):2841–2852.

27. Bonvalot L, Mercury M, Zerega Y: Laboratory-scale apparatus for the characterization of gas-phase dioxin adsorption. Instrumentation Science & Technology 2016:1–15.

28. Xuan Wan PWS: Volatile Sex Pheromone Extraction and Chemoattraction Assay in *Caenorhabditis elegans*.. JoVE (Journal of Visualized Experiments) 2024, 210 (2024): e67115.

29. Ding K HY, Seid TW, Buser C, Karigo T, Zhang S, Dickman DK, Anderson DJ.: Imaging neuropeptide release at synapses with a genetically engineered reporter. Elife 2019, 8:e46421.

30. Jurenka RA, Haynes KF, Adlof RO, Bengtsson M, Roelofs WL: Sex pheromone component ratio in the cabbage looper moth altered by a mutation affecting the fatty acid chain-shortening reactions in the pheromone biosynthetic pathway. Insect biochemistry and molecular biology 1994, 24(4):373–381.

31. Rasmussen LE, Lee TD, Zhang A, Roelofs WL, Daves GD, Jr.: Purification, identification, concentration and bioactivity of (Z)-7-dodecen-1-yl acetate: sex pheromone of the female Asian elephant, Elephas maximus. Chemical senses 1997, 22(4):417–437.

32. Naka H, Nakazawa T, Sugie M, Yamamoto M, Horie Y, Wakasugi R, Arita Y, Sugie H, Tsuchida K, Ando T: Synthesis and characterization of 3,13- and 2,13-octadecadienyl compounds for identification of the sex pheromone secreted by a clearwing moth, Nokona pernix. Biosci Biotechnol Biochem 2006, 70(2):508–516.

33. Khallaf MA, Cui R, Weißflog J, Erdogmus M, Svatoš A, Dweck HK, Valenzano DR, Hansson BS, Knaden M: Large-scale characterization of sex pheromone communication systems in Drosophila. Nature communications 2021, 12(1):4165.

34. Gomez-Diaz C, Benton R: The joy of sex pheromones. EMBO reports 2013, 14(10):874–883.

35. Yoshida K, Hirotsu T, Tagawa T, Oda S, Wakabayashi T, Iino Y, Ishihara T: Odour concentration-dependent olfactory preference change in C. elegans. Nature communications 2012, 3(1):739.

36. Taniguchi G, Uozumi T, Kiriyama K, Kamizaki T, Hirotsu T: Screening of odor-receptor pairs in *Caenorhabditis elegans* reveals different receptors for high and low odor concentrations. Science signaling 2014, 7(323):ra39-ra39.

37. Luo L, Gabel CV, Ha H-I, Zhang Y, Samuel AD: Olfactory behavior of swimming *C. elegans* analyzed by measuring motile responses to temporal variations of odorants. Journal of neurophysiology 2008, 99(5):2617–2625.

38. Bargmann CI: Chemosensation in C. elegans. WormBook: the online review of Celegans biology 2006:1-29.

39. Lanjuin A, VanHoven MK, Bargmann CI, Thompson JK, Sengupta P: Otx/otd homeobox genes specify distinct sensory neuron identities in C. elegans. Dev Cell 2003, 5(4):621–633.

40. Ikeda M, Nakano S, Giles AC, Xu L, Costa WS, Gottschalk A, Mori I: Context-dependent operation of neural circuits underlies a navigation behavior in Caenorhabditis elegans. Proc Natl Acad Sci U S A 2020, 117(11):6178–6188.

41. Sengupta P, Colbert HA, Bargmann CI: The *C. elegans* gene odr-7 encodes an olfactory-specific member of the nuclear receptor superfamily. Cell 1994, 79(6):971–980.

42. Ferguson EL, Horvitz HR: Identification and characterization of 22 genes that affect the vulval cell lineages of the nematode *Caenorhabditis elegans*. Genetics 1985, 110(1):17–72.

43. Schwartz HT, Horvitz HR: The *C. elegans* protein CEH-30 protects male-specific neurons from apoptosis independently of the Bcl-2 homolog CED-9. Genes Dev 2007, 21(23):3181–3194.

44. Troemel ER, Sagasti A, Bargmann CI: Lateral signaling mediated by axon contact and calcium entry regulates asymmetric odorant receptor expression in *C. elegans*. Cell 1999, 99(4):387–398.

45. Bauer Huang SL, Saheki Y, Vanhoven MK, Torayama I, Ishihara T, Katsura I, Van Der Linden A, Sengupta P, Bargmann CI: Left-right olfactory asymmetry results from antagonistic functions of voltage-activated calcium channels and the Raw repeat protein OLRN-1 in *C. elegans*. Neural Development 2007, 2(1):24.

46. Werner KM, Perez LJ, Ghosh R, Semmelhack MF, Bassler BL: *Caenorhabditis elegans* recognizes a bacterial quorum-sensing signal molecule through the AWCON neuron. J Biol Chem 2014, 289(38):26566–26573.

47. Honzel BE, Foley SJ, Politz SM: *C. elegans srf-6* and *nsy-1* mutations result in a similar 2AWC^ON^ phenotype and do not complement (*srf-6* is *nsy-1* II). MicroPublication Biology 2019, 2019:10.17912/micropub.biology.000128.

48. Kotera I, Tran NA, Fu D, Kim JH, Byrne Rodgers J, Ryu WS: Pan-neuronal screening in *Caenorhabditis elegans* reveals asymmetric dynamics of AWC neurons is critical for thermal avoidance behavior. eLife 2016, 5.

49. Tian L, Hires SA, Mao T, Huber D, Chiappe ME, Chalasani SH, Petreanu L, Akerboom J, Mckinney SA, Schreiter ER et al: Imaging neural activity in worms, flies and mice with improved GCaMP calcium indicators. Nature Methods 2009, 6(12):875–881.

50. Chalasani SH, Chronis N, Tsunozaki M, Gray JM, Ramot D, Goodman MB, Bargmann CI: Dissecting a circuit for olfactory behaviour in Caenorhabditis elegans. Nature 2007, 450(7166):63-70.

51. Hsueh YP, Gronquist MR, Schwarz EM, Nath RD, Lee CH, Gharib S, Schroeder FC, Sternberg PW: Nematophagous fungus Arthrobotrys oligospora mimics olfactory cues of sex and food to lure its nematode prey. Elife 2017, 6.

52. Von Reuss SH, Schroeder FC: Combinatorial chemistry in nematodes: modular assembly of primary metabolism-derived building blocks. Natural Product Reports 2015, 32(7):994–1006.

